# Alterations and Imbalance of Dorsal and Ventral Mossy Cells in a Mouse Model of Epilepsy

**DOI:** 10.1101/2024.11.07.622490

**Authors:** Christine S. Huang, Carolyn R. Houser

**Author notes:** Correspondence should be addressed to Carolyn R. Houser at. This study is dedicated to the memory of Dr. Zechun Peng who was instrumental in planning and conducting the experiments described in this paper. The authors declare no competing financial interests.

## Abstract

Mossy cells (MCs) in the hilus of the dentate gyrus (DG) are important for regulating activity of dentate granule cells and are particularly vulnerable to excitotoxic damage in epilepsy. Recent studies have demonstrated that MCs in the dorsal and ventral DG differ in the patterns of their axonal projections and neurochemical identities. Such differences raised questions about the vulnerability and plasticity of dorsal and ventral MCs in epilepsy and led to this study using a mouse pilocarpine model of epilepsy. Dorsal MCs were labeled by transfection of Cre-dependent eYFP in the dorsal DG of *Calcrl-Cre* mice that express Cre selectively in MCs. Ventral MCs were labeled with calretinin (CR), which labels ventral but not dorsal MCs. At 6-8 weeks after pilocarpine treatment, MC loss and axonal projections of remaining MCs were studied in control and pilocarpine-treated mice with confocal microscopy. Dorsal MCs were severely depleted, but many ventral MCs remained, and quantitative analysis of GluA2-labeled hilar neurons demonstrated a proportionally greater loss of dorsal MCs (77.6% loss) than ventral MCs (21.5% loss). Loss of dorsal MCs led to a marked reduction in the dorsal commissural pathway, while the remaining ventral MCs maintained a prominent, though reduced, ventral to dorsal association pathway. In pilocarpine-treated animals, a plexus of CR-labeled fibers extended into the middle molecular layer, suggesting axonal sprouting of remaining ventral MCs, with some of these fibers in contact with parvalbumin-labeled dendrites. These findings suggest that dorsal and ventral MCs differ in their vulnerability to seizure-induced damage in this animal model, creating an imbalance between the dorsal and ventral MC pathways that could alter the excitatory/inhibitory balance within the dentate gyrus.

## Introduction

Mossy cells (MCs) are a major group of glutamatergic neurons within the dentate gyrus (DG) that are important for regulating activity of dentate granule cells (Scharfman, 2016). By maintaining sparse activity of dentate granule cells, MCs contribute to the processes of learning and memory, including pattern separation, and may also control excitability within the DG (Jinde et al., 2012; Bui et al., 2018; GoodSmith et al., 2019; Galloni et al., 2022; Huang et al., 2024). Determining the specific roles of MCs has remained challenging, due to their unique circuitry and multiple pathways. MCs form excitatory contacts with both granule cells and interneurons (Scharfman, 1995; Wenzel et al., 1997) and, through these connections, can exert either direct excitatory or indirect inhibitory effects on granule cells (Buzsàki and Eidelberg, 1981; Jinde et al., 2012; Hsu et al., 2016; Abdulmajeed et al., 2022). In addition, MCs form both local commissural pathways and extensive association pathways that extend longitudinally through the DG. These axonal projections are critical for associating activity in one region of the DG with that in distant regions (Amaral and Witter, 1989; Buckmaster and Schwartzkroin, 1994). However, in pathological conditions, the association fibers could allow excessive activity at one level of the DG to be propagated over long distances and thus increase excitability throughout the DG (Buckmaster et al., 1996; Bui et al., 2018).

MCs are also among the most vulnerable neurons in the DG, and loss of MCs has been observed in animal models of traumatic brain injury (TBI) and acquired epilepsy, as well as human temporal lobe epilepsy (Lowenstein et al., 1992; Blümcke et al., 2000; Sloviter et al., 2003; Jiao and Nadler, 2007). Despite their vulnerability, a considerable number of MCs often remain (Santhakumar et al., 2000). This has led to questions about their role in epilepsy and whether it is the loss of MCs and their connections to inhibitory interneurons or excessive activity of remaining MCs and their innervation of granule cells that could contribute to a pro-epileptic state.

MCs are present throughout the DG, and it had been assumed that all MCs had similar patterns of termination in the inner molecular layer of the DG. However, recent studies have demonstrated that dorsal and ventral MCs have different patterns of axonal projections (Botterill et al., 2021; Houser et al., 2021). While ventral MCs exhibit the classical pattern and densely innervate the inner molecular layer, dorsal MCs form a band of commissural fibers at dorsal levels but form a more diffuse projection in the middle molecular layer at more ventral levels (Houser et al., 2021). These different patterns, along with other differences in the neurochemical, physiological and morphological features of MCs (Liu et al., 1996; Jinno et al., 2003; Bienkowski et al., 2018), suggest that dorsal and ventral MCs may have different functional roles. Recent behavioral studies are now demonstrating such differences (Fredes et al., 2021; Wang et al., 2021; Jeong et al., 2024).

Recognition of differences in dorsal and ventral MCs and their axonal projections led to questions about possible differences in the extent of MC loss and plasticity in these two groups of MCs in a model of acquired epilepsy. The major goals of this study were to 1) compare the loss of dorsal and ventral MCs in a mouse pilocarpine model of epilepsy during the chronic stage; 2) compare the resulting patterns of axonal projections of each group of MCs; and 3) determine if remaining MCs exhibit axonal reorganization.

## Materials and Methods

### Animals

Two mouse lines were used in this study of MCs in control and pilocarpine-treated mice -- the *Calcrl-Cre* transgenic mouse that expresses Cre relatively selectively in MCs throughout the DG and C57BL/6J mice. The *Calcrl-Cre* mouse expresses Cre under the control of the murine calcitonin receptor-like promoter and was generated and characterized by Dr. Kazu Nakazawa and colleagues (Jinde et al., 2012). Material for re-derivation of the *Calcrl-Cre* mice was generously provided by Dr. Nakazawa, and a breeding colony was established. These mice are now available from Jackson Laboratory (C57BL/6N-Tg(Calcrl,cre)4688Nkza/J, JAX stock #023014). C57BL/6J mice were obtained from Jackson Laboratory (JAX stock #000664).

The *Calcrl-Cre* mice were essential for identifying dorsal MCs and their fibers as there is currently no neurochemical marker for selectively labeling this group of MCs. Dorsal MCs were thus identified in these mice following transfection for eYFP restricted to the dorsal DG. Immunohistochemical labeling for CR was used to identify ventral MCs and their fibers in both *Calcrl-Cre* and C57BL/6J mice.

Mice were 2-3 months of age at the time of pilocarpine or control treatment and were prepared for neuroanatomical study at 4-5 months of age (*Calcrl-Cre* mice: n = 5 pilocarpine-treated and 4 controls; C57BL/6J mice: n = 3 pilocarpine-treated and 3 controls). All animal use protocols conformed to the National Institutes of Health guidelines and were approved by the University of California, Los Angeles, Chancellor’s Animal Research Committee.

### Pilocarpine treatment

The pilocarpine mouse model of recurrent seizures has been described in detail in previous studies (Peng et al., 2004; Peng et al., 2013). Briefly, a single episode of seizure activity (status epilepticus) is induced; the animals progressively recover over the next 1-2 weeks; and they then begin exhibiting spontaneous behavioral seizures, characteristic of epilepsy. In the current study, mice were first injected with scopolamine methyl nitrate (1 mg/kg, s.c.; Sigma-Aldrich) to reduce peripheral cholinergic effects. Thirty min later, experimental animals (both *Calcrl-Cre* and C57BL/6J mice) received an injection of pilocarpine hydrochloride (310-340 mg/kg, s.c.; Sigma-Aldrich) to induce status epilepticus. At 2 h after the onset of sustained seizures, diazepam (7-10 mg/kg, s.c.; Hospira) was administered to suppress or ameliorate the acute behavioral seizures. Control animals received an identical series of injections except that pilocarpine was replaced with a similar volume of sterile saline. Following the induced seizure episode, pilocarpine-treated animals recovered and resumed normal behavior over the next few days. At 3 weeks following pilocarpine treatment, the *Calcrl-Cre* mice received a unilateral transfection of an adeno-associated viral (AAV) vector to label MCs with eYFP.

### Viral vector injections

To selectively label dorsal MCs, *Calcrl-Cre* mice were transfected with a Cre-dependent double-floxed recombinant AAV vector containing a construct encoding eYFP in the hilus of the dorsal DG. The viral vector for eYFP (AAV DJ-EF1a-DIO eYFP) was obtained from the Neuroscience Gene Vector and Virus Core, Stanford University.

For transfections, mice were anesthetized with isoflurane, and a Cre-dependent viral vector was stereotaxically injected in the dorsal DG with a Nanoject II injector (Drummond Scientific), using glass pipettes. A small animal stereotaxic instrument with digital display console (Model 940; Kopf Instruments) was used for precise positioning of the pipette in the hilus. Injections for eYFP were made unilaterally in the dorsal hilus (right side). The goal was to label the rostral group of MCs as completely as possible and avoid labeling in the middle or caudal regions of the DG. Optimal stereotaxic coordinates were determined previously (Houser et al., 2021) and were made at two sites in close proximity to each other (−1.8 anteroposterior (AP), 1.0 mediolateral (ML), 2.2 dorsoventral (DV); and -2.0 AP, 1.2 ML, 2.1 DV) (Paxinos and Franklin, 2019). Injection volumes were 69-92 nl at each site (23 nl increments x 3-4 injections). Following each injection, the pipette was left in position for 5 min before it was slowly retracted from the brain. At 3 weeks post transfection, the *Calcrl-Cre* mice were prepared for neuroanatomical studies.

### Tissue preparation for light microscopy

All mice were deeply anesthetized with Fatal-Plus (90 mg/kg, i.p.) and perfused transcardially with 4% paraformaldehyde in 0.12 M phosphate buffer (pH 7.3). After 1 h in situ at 4°C, brains were removed and postfixed for 1 h; rinsed and cryoprotected in a 30% sucrose solution overnight; embedded in OCT compound (Sakura Finetek) and frozen on dry ice; and sectioned at 30 μm with a cryostat (CM 3050S, Leica Microsystems). For detailed comparisons of MCs and their projections at dorsal and ventral levels, brains were sectioned coronally through the dorsal (rostral) third of the hippocampus and horizontally through the ventral (caudal) two-thirds of the hippocampus.

### Antisera

Dorsal MCs and their fibers were distinguished by their expression of transfected eYFP, as there is currently no selective immunohistochemical marker for these neurons. All eYFP labeling was from the transfection-induced expression of the fluorescent protein and was not enhanced by immunofluorescent labeling. However, immunohistochemical studies were used to identify the entire population of MCs for cell counts, ventral MCs, and PV interneurons. A rabbit polyclonal antiserum to GluA2 (Millipore, AB1768), an AMPA receptor subunit (previously referred to as GluR2), was used as a general marker for MCs within the hilus (Leranth et al., 1996; Petralia et al., 1997; Jiao and Nadler, 2007). A rabbit polyclonal antiserum to CR (Swant, Cat. #7697) was used to selectively label ventral MCs and their projections (Liu et al., 1996; Blasco-Ibanez and Freund, 1997). A mouse monoclonal antibody to PV (Millipore, MAB1572) was used to identify PV+ interneurons and determine their relationship to MC axons.

### Immunoperoxidase labeling

For broad labeling of all MCs in the hilus and associated MC counts, GluA2 was localized with standard avidin-biotin-peroxidase methods. Free-floating sections were incubated in 1% H2O2 for 30 min to reduce endogenous peroxidase-like activity. After rinsing in 0.1M Tris buffered saline (TBS; pH 7.3), sections were incubated in 10% normal goat serum diluted in TBS containing 0.3% Triton X-100 for 2-3 h to reduce non-specific binding and increase penetration of the antibodies. Sections were incubated in primary GluA2 antiserum (1:100) overnight at room temperature; incubated in biotinylated secondary antiserum (goat anti-rabbit IgG, 1:200; Vector Laboratories) at room temperature for 1 h; and then incubated in avidin-biotin peroxidase complex (1:100, Vectastain Elite ABC kit; Vector Laboratories) for 1 h. To visualize the peroxidase labeling, sections were processed with Stable diaminobenzidine (DAB) (Invitrogen) for 12 min, and immunolabeling was enhanced by incubation in 0.003% osmium tetroxide in phosphate-buffered saline (PBS) for 30 seconds. Sections were mounted on slides, dehydrated and coverslipped.

### Immunofluorescence labeling

For double-labeling studies, CR was localized with immunofluorescence methods in sections from animals previously transfected for eYFP in order to distinguish between dorsal (eYFP-labeled) and ventral (CR-labeled) mossy cells in the same sections. In addition, PV was localized in animals previously transfected for eYFP to determine the relationship between eYFP-labeled MC fibers and PV+ interneurons. For both sets of studies, sections were incubated for 2 h in 10% normal goat serum to block nonspecific binding sites and 0.3% Triton X-100. Sections were incubated in primary antiserum (CR, 1:2K or PV, 1:5K) for 72 h. Sections were then incubated in secondary antiserum (goat anti-rabbit or goat anti-mouse IgG conjugated to Alexa Fluor 555; Invitrogen) for 4 h at room temperature, mounted on slides, and coverslipped with antifade medium ProLong Diamond (Invitrogen). In an additional set of studies, CR and PV were localized with double immunofluorescence methods to determine the relationship of CR-labeled fibers and PV+ interneurons. Sections were pretreated as described above and then incubated in a cocktail of antisera (CR, 1:2K and PV, 1:5K for 72 h. Sections were then incubated in secondary antisera (goat anti-rabbit conjugated to Alexa Fluor 555 and goat anti-mouse conjugated to Alexa Fluor 488; Invitrogen) for 4h, and processed as described above.

### Cell counting methods and analysis

For MC counting, the hilar region was delineated by tracing along the inner border of the granule cell layer and drawing straight lines from each end of the granule cell layer to the proximal tip of the CA3 pyramidal cell layer. This method defined the hilus reliably among sections, and virtually all putative MCs were included within the region, even when the outlined region did not precisely match the anatomical borders of the hilus, as in some horizontal sections. MCs were identified as large (>10 um somal diameter) GluA2-labeled neurons with a clear nucleus. These parameters excluded ectopic granule cells that could potentially be increased in the pilocarpine-treated animals.

GluA2+ MCs were counted in the hilus on each side at 3 levels of the dorsal DG in coronal sections (300 μm intervals) and 3 levels of the ventral DG in horizontal sections (500 μm intervals) in control and pilocarpine-treated animals. Data from the 3 levels of the dorsal and ventral dentate gyrus, respectively, were combined and expressed as the mean number of GluA2 cells/hilus in the dorsal and ventral regions. Statistical calculations and analyses were done in GraphPad Prism 10 (GraphPad) and Excel 365, using two-tailed Mann-Whitney tests. Significance level was set at *p* < 0.05.

### Analysis of immunofluorescence data

Fluorescence-labeled sections were scanned with an LSM 880 (Carl Zeiss) confocal microscope, and confocal images were analyzed with Zen 3.6 software (Blue edition, Carl Zeiss). For detailed analyses of eYFP- and CR-labeled MCs in the hilus and their axonal projections throughout the DG, maximum intensity Z-stack projection images were acquired with excitation spectra 488 and 555 and optical slice thicknesses set at 0.5-2 μm. In sections with two fluorescent labels, optical slices were scanned separately for each label, alternating between the two channels in each slice through the Z-stack. Images were acquired with Plan-Apochromat objectives (20X air, 40X water, 63X oil; Carl Zeiss). Whenever possible, sections from control and pilocarpine-treated animals from the same immunofluorescence experiments were imaged with the same conditions.

## Results

A major goal of this study was to compare the patterns of cell loss in dorsal and ventral groups of MCs, using a specific marker for each group. Dorsal MCs were identified by their expression of eYFP following unilateral transfection of Cre-dependent eYFP in the dorsal one-third of the DG in *Calcrl-Cre* mice in which Cre is selectively expressed in MCs (Jinde et al., 2012). Ventral MCs were identified by immunohistochemical localization of CR that labels ventral but not dorsal MCs in the mouse (Liu et al., 1996; Blasco-Ibanez and Freund, 1997; Fujise et al., 1998).

### eYFP-labeled MCs in the dorsal DG were severely reduced in pilocarpine-treated mice

In control mice, eYFP-labeled MCs in the dorsal DG were abundant throughout the hilus on the transfected side (Fig. 1B), replicating previous findings in these mice (Houser et al., 2021). These MCs form the commissural pathway of the dorsal DG, and a dense band of eYFP-labeled fibers was evident in the inner molecular layer, slightly above the granule cell layer, on the contralateral side (Fig. 1A). A more diffuse plexus of eYFP-labeled fibers extended beyond the dense band of fibers and into the middle molecular layer (Fig. 1A). Labeled fibers were also evident in the hilus (Fig. 1A).

**Figure 1.**
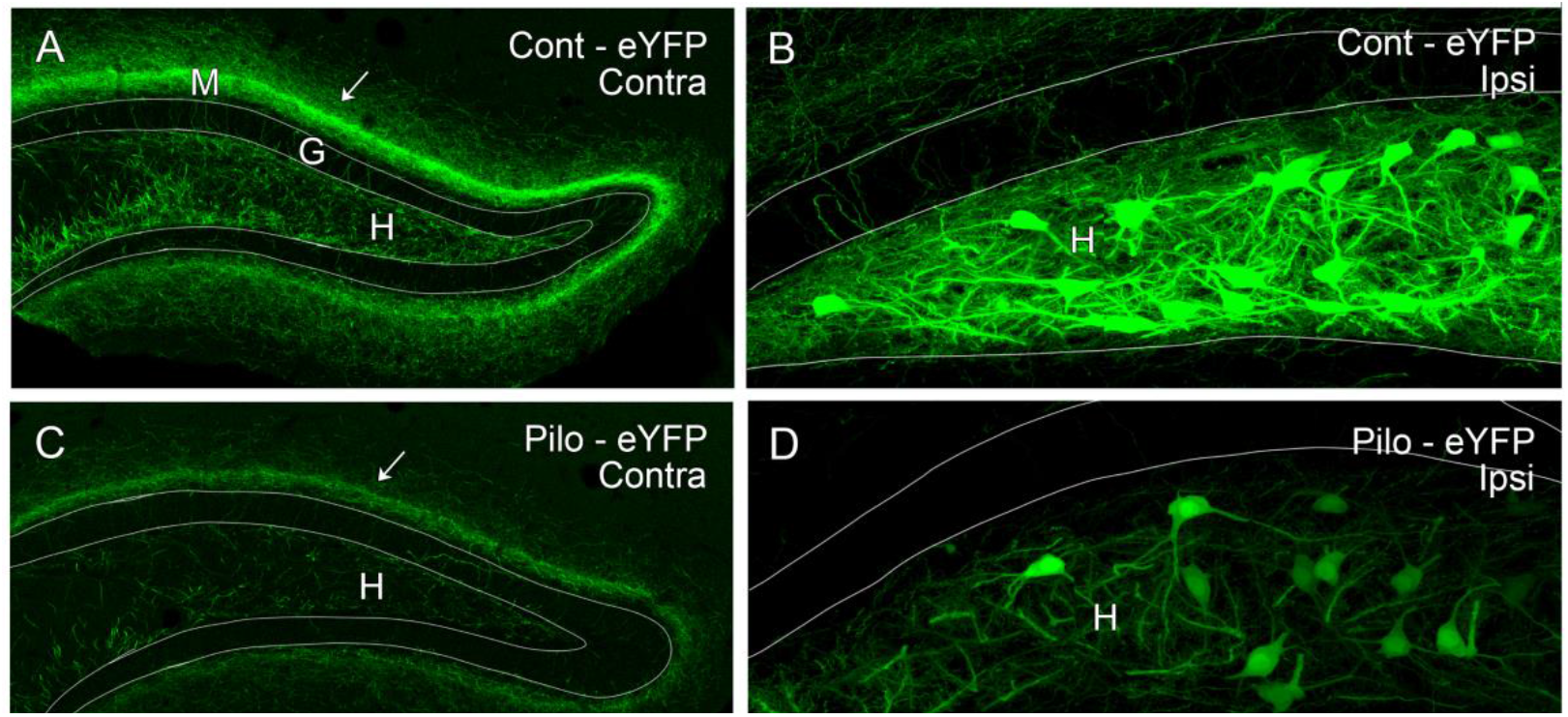
eYFP-labeled dorsal MCs and their commissural axonal projections are abundant in a control mouse (**A**,**B**) but are severely depleted in a pilocarpine (Pilo)-treated mouse (**C**,**D**). **A**,**B**, In a control mouse, numerous MCs fill the hilus (H) following unilateral transfection of eYFP in the dorsal DG of a *Calcrl-Cre* mouse (**B**). Commissural fibers from these neurons form a prominent band in the molecular layer (M) on the contralateral side (**A**). A dense portion of the band is located at a slight distance above the granule cell layer (G), and a more diffuse plexus of fibers extends into the middle molecular layer (arrow). **C**,**D**, A similar transfection in a pilo-treated mouse labels the few remaining MCs within the hilus (**D**), and the band of commissural fibers on the contralateral side is substantially reduced (arrow) (**C**). Scale bars: 100 μm (**A**,**C**); 25 μm (**B**,**D**).

In the pilocarpine-treated animals, eYFP-labeled MCs were substantially reduced in the dorsal DG (Fig. 1D) compared to those in the control animals (Fig. 1B). The band of eYFP-labeled commissural fibers at the same rostral-caudal level of the DG was severely reduced in width and density (Fig. 1C), consistent with the sparsity of dorsal MCs on the opposite side. Within the dorsal DG, the extent of MC loss was most striking at more rostral levels, where MC loss was often nearly complete (Fig. 2A,B). The remaining isolated cells exhibited a typical MC morphology (Fig. 2A).

**Figure 2.**
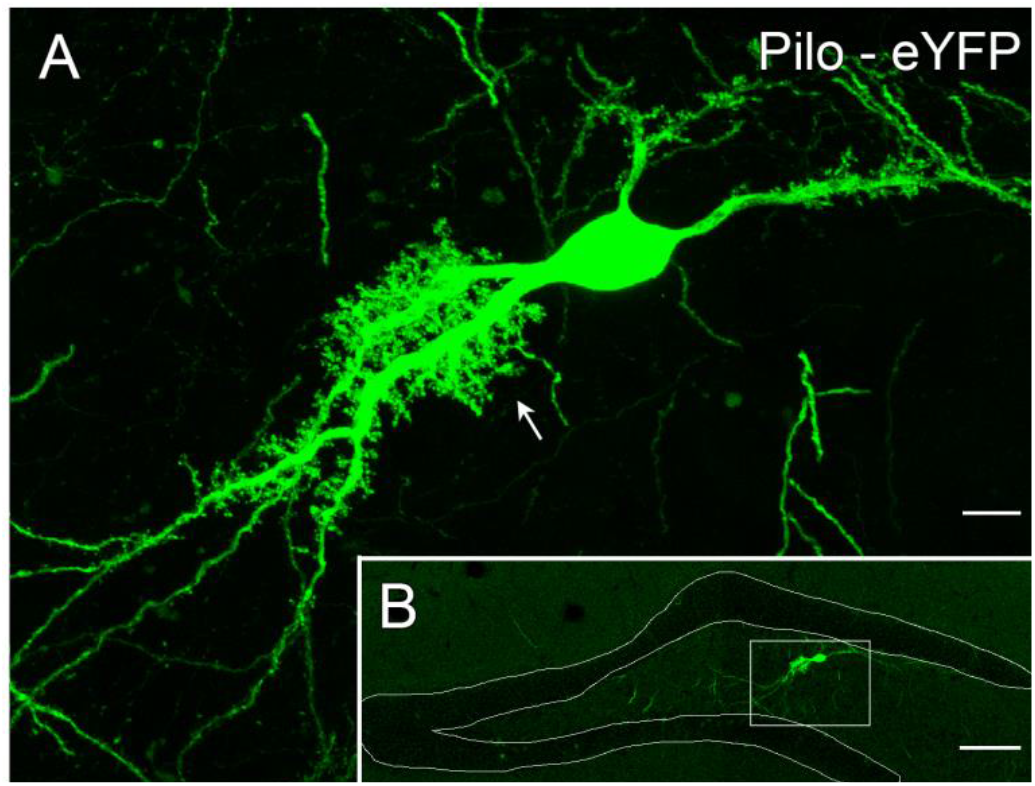
A single MC is evident in a section through the dorsal hilus of a pilo-treated animal. **A**,**B**, In an animal with severe cell loss in the dorsal DG, an isolated eYFP-labeled cell remains within the section (**B**) and exhibits a typical MC morphology, with a high concentration of spines on the proximal dendrites (arrow) (**A**). Scale bars: 10 μm (**A**); 100 μm (**B**).

This qualitative analysis demonstrated a substantial loss of dorsal MCs and a related reduction of their commissural fibers in the pilocarpine-treated mouse.

### CR-labeled MCs of the ventral DG were moderately reduced in pilocarpine-treated mice

CR immunohistochemistry was used to label the cell bodies of ventral MCs and their associated axonal projections. In control animals, the ventral MC somata were labeled throughout the hilus and clearly delineated this region (Fig. 3A-C). In pilocarpine-treated animals, the labeled MCs were distributed more diffusely and appeared to be reduced in number (Fig. 3D-F). In 6 of 8 pilocarpine-treated animals, MCs appeared most severely depleted in the medial region of the hilus in horizonal sections (Fig. 3D-F). The band of MC fibers in the inner molecular layer was present in all animals but the density of the fibers was reduced in the pilocarpine-treated mice, consistent with some loss of MCs (compare Fig. 3A-C with D-F). This qualitative analysis of CR-labeled MCs suggested a mild to moderate loss of ventral MCs and their axonal association fibers in the inner molecular layer.

**Figure 3.**
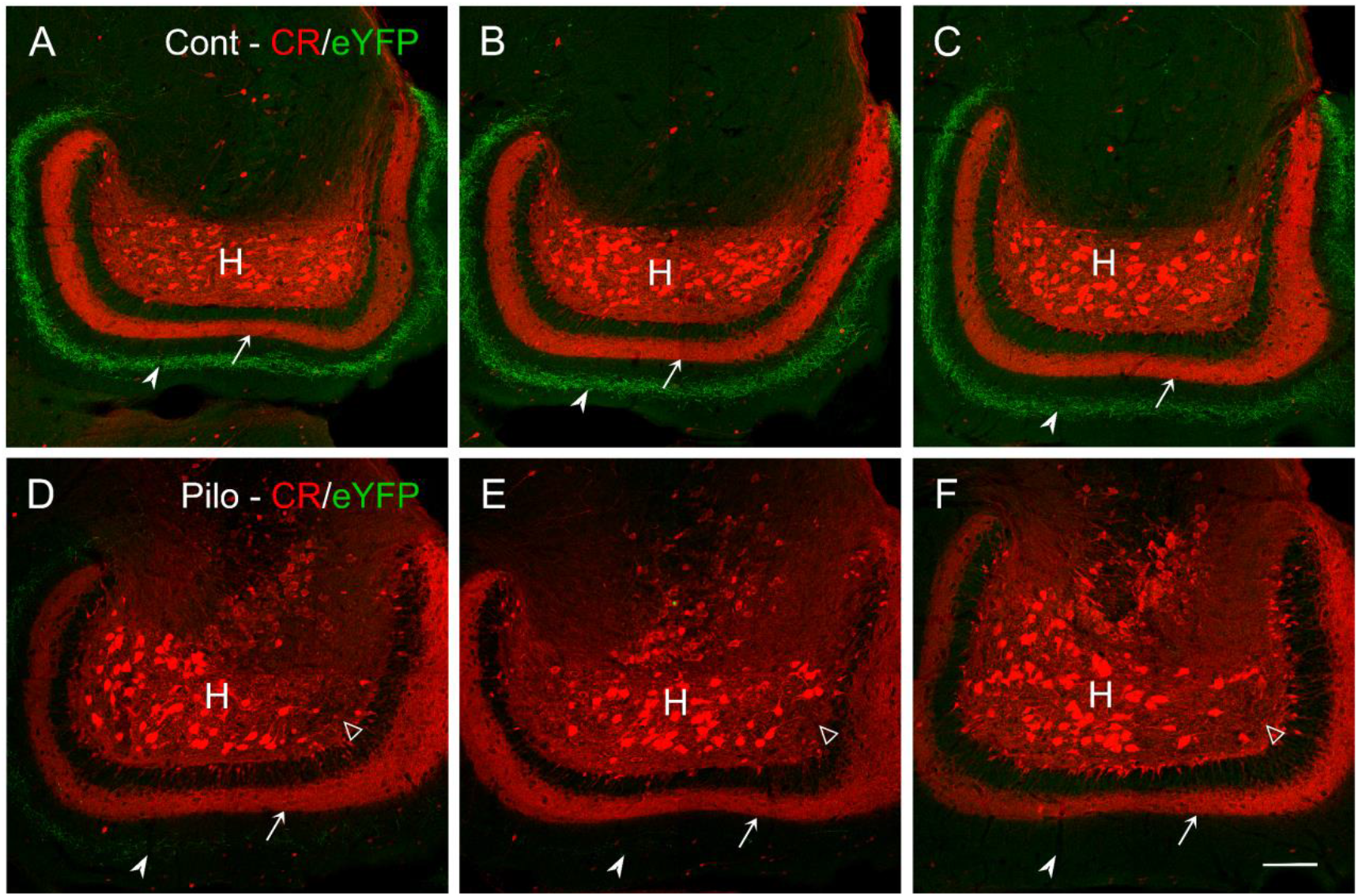
Numerous calretinin (CR)-labeled MCs fill the ventral hilus of the DG in control mice but appear fewer and more dispersed in pilo-treated mice. **A-C**, MCs delineate the ventral hilus (H) in three control mice, and a prominent band of ventral MC fibers is present in the inner molecular layer (arrows). A more diffuse band of eYFP-labeled fibers from dorsal MCs is evident in the middle molecular layer (arrowheads). **D-F**, MCs appear less dense in the ventral hilus of three pilo-treated mice, with the most obvious loss in the medial region of the hilus (open arrowheads). A band of ventral MC fibers remains in the inner molecular layer (arrows) but appears less dense than in control animals. The band of dorsal MC fibers, normally present in the middle molecular layer (see **A-C**), is severely reduced or absent (arrowheads). Scale bars: 100 μm (**A-F**).

### Dorsal MCs were more severely depleted than ventral MCs in pilocarpine-treated mice

For quantitative analyses and comparisons of cell loss in the dorsal and ventral DG, coronal sections of the dorsal and horizontal sections of the ventral hippocampus were labeled for GluA2 which serves as a general marker of MCs. This allowed comparisons of similarly labeled MCs in both the dorsal and ventral hilus.

In coronal sections, a distinct loss of GluA2-labeled MCs was evident at all levels (compare Fig. 4A,B and 4E,F). However, the GluA2 cell loss in the dorsal DG was greater at more rostral levels, where the loss was often nearly complete (Fig. 4E). At ventral levels of the DG, cell loss was evident but was less striking than at dorsal levels (compare Fig. 4C,D & G,H). These findings were consistent with our initial findings of a marked loss of eYFP-labeled MCs in the dorsal DG and a potentially more modest loss of CR-labeled MCs in the ventral DG.

**Figure 4.**
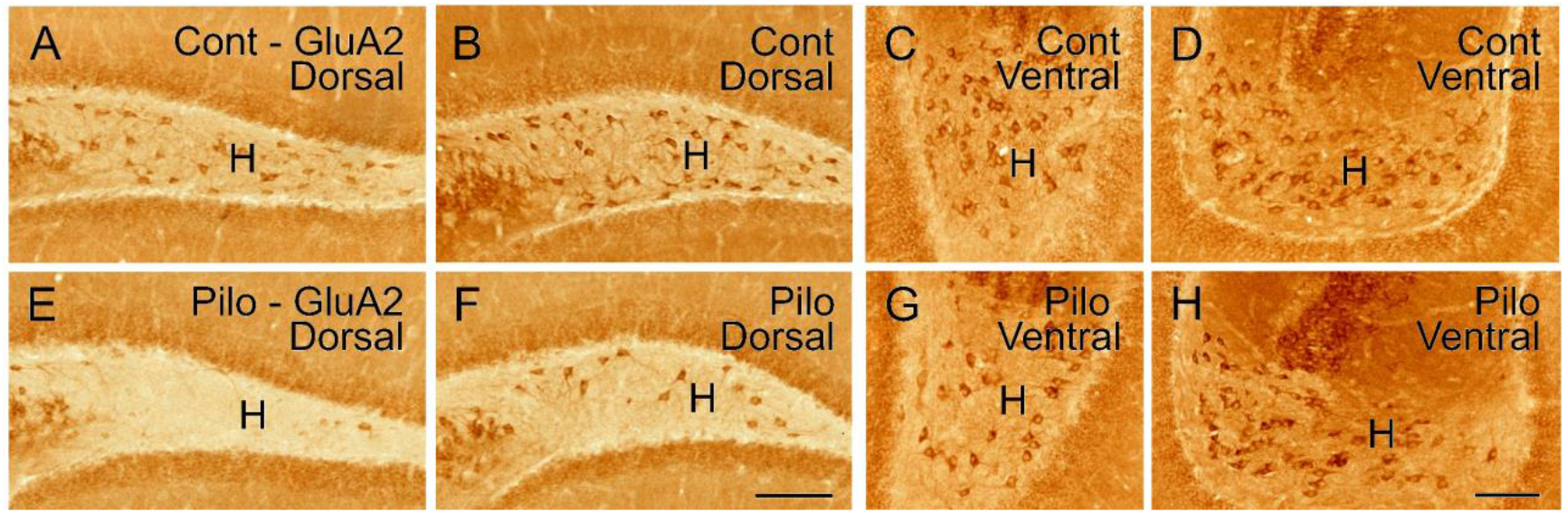
GluA2-labeled MCs are abundant in the dorsal and ventral DG of control (Cont) animals (**A-D**) but appear reduced at similar levels in pilo-treated animals (**E-H**). **A**,**B & E**,**F**, In coronal sections at two levels of the dorsal DG, GluA2-labeled neurons are dispersed throughout the hilus (H) in control animals (**A**,**B**) but are severely depleted in sections at the same levels in pilo-treated animals (**E**,**F**). **C**,**D & G**,**H**, In horizontal sections at two levels of the ventral DG in control animals, GluA2-labeled neurons are relatively evenly distributed throughout the hilus (**C**,**D**). In pilo-treated animals, GluA2-labeled neurons appear less numerous than in controls (**G-H**), but the differences are less striking than at dorsal levels (compare with **E**,**F**). Scale bars: 100 μm (**A**,**B**,**E**,**F**); 50 μm (**C**,**D**,**G**,**H**).

For quantitative analyses, GluA2+ MCs were counted in the hilus on each side at 3 levels of both dorsal and ventral DG in control and pilocarpine-treated animals. Data from the 3 levels of the dorsal and ventral DG, respectively, were combined and expressed as the mean number of GluA2 cells/hilus in each region (N = 6 hilar regions from both dorsal and ventral regions x 7-8 control and 7-8 pilocarpine treated animals = 42-48 hilar regions for dorsal and ventral DG respectively).

In dorsal sections, greater numbers of GluA2+ neurons per hilus were found in control mice (26.2 +/-1.0 cells) than in pilocarpine-treated-mice (5.8 +/-0.8 cells; p<0.001, Mann-Whitney) (Fig. 5A). In ventral sections, greater numbers of GluA2+ neurons per hilus were also found in control mice (66.8 +/-4.0 cells) than in pilocarpine-treated mice (52.5 +/-5.7 cells; p<0.03, Mann-Whitney), but the differences were less (Fig. 5A). Based on these data, the percentages of cells remaining in the pilocarpine-treated animals were 22.4% in the dorsal DG and 78.5% in the ventral DG (Fig. 5B), and the percentages of cells remaining in the dorsal and ventral DG in pilocarpine-treated animals were significantly different (p<0.001) (Fig. 5B). The resulting percentage cell loss in the pilocarpine-treated animals was 77.6% in the dorsal hilus and 21.5% in the ventral hilus.

**Figure 5.**
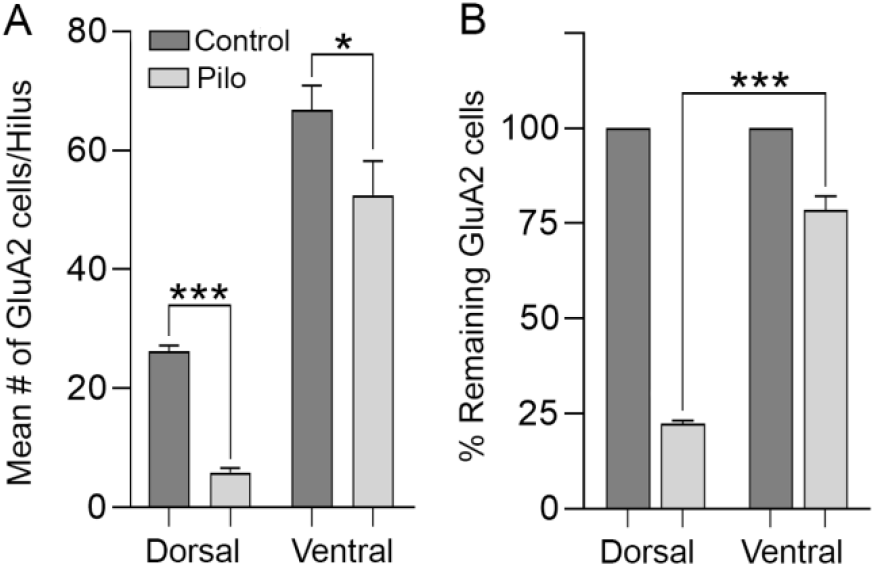
Quantitative analyses demonstrated significant differences in the number of GluA2-labeled MCs remaining in control and pilo-treated animals. **A**, The number of GluA2 cells/hilus in the dorsal hilus was reduced from a mean of 26.2 cells/hilus in controls to 5.8 cells/hilus in pilo-treated animals (p<0.001 (***). The number of cells in the ventral hilus was also reduced, but to a lesser extent, from a mean of 66.8 cells/hilus in controls to 52.5 cells/hilus in pilo-treated animals (p<0.03). **B**, The percentage of cells remaining in pilo-treated animals compared to controls was 22.4% in the dorsal hilus and 78.5% in the ventral hilus. Data were obtained from 6 hilar regions at both dorsal and ventral levels in each of 7-8 control and 7-8 pilo-treated animals.

The loss of GluA2-labeled MCs was also compared at each of the 6 levels of the DG that were used for the cell counts, from most dorsal to most ventral. The mean number of GluA2+ neurons per hilus was determined for each level in 6-8 control and 7-8 pilocarpine-treated mice. In control mice, GluA2+ neurons were more numerous in the ventral hilus and least numerous in the dorsal hilus (Fig. 6), as described previously in rats and mice (Jiao and Nadler, 2007; Volz et al., 2011). However, the differences in cell numbers between control and pilocarpine-treated animals were greatest at dorsal levels and became progressively less at ventral levels, where many GluA2+ cells remained in the pilocarpine-treated mice (Fig. 6).

**Figure 6.**
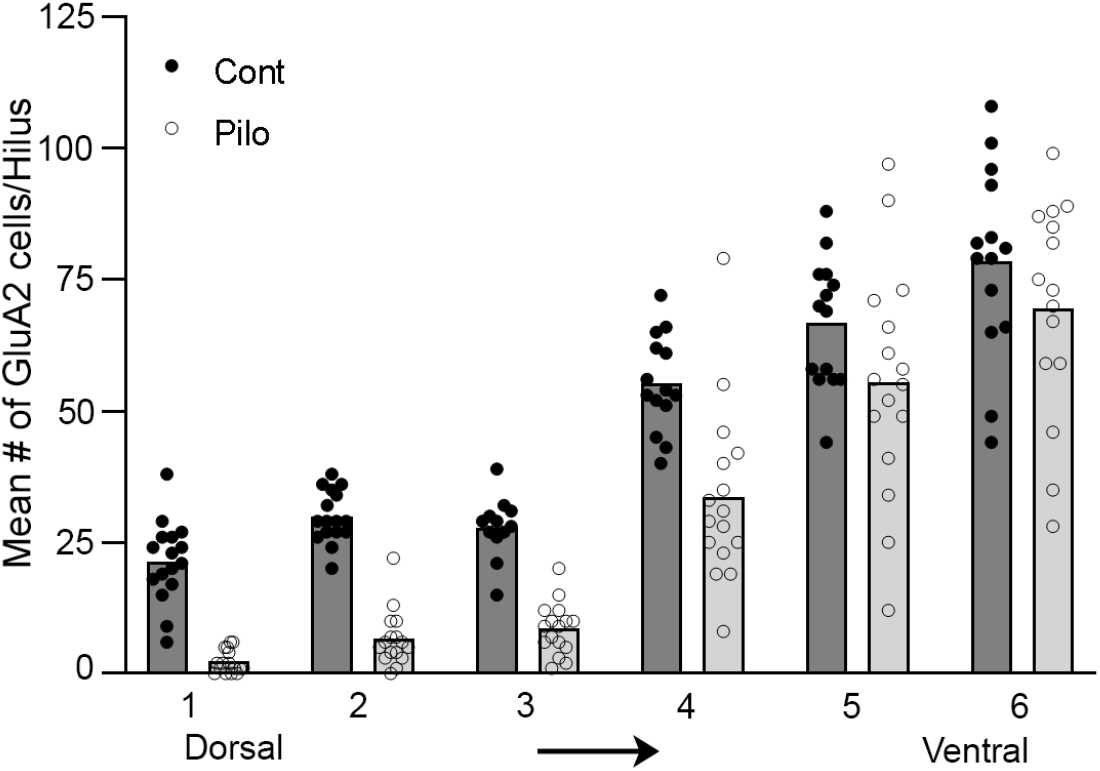
Comparison of the mean number of GluA2-labeled cells/hilus at 3 levels of the dorsal and 3 levels of the ventral DG in control and pilo-treated animals. Each circle represents the number of cells in a single hilus (2 hilar regions per animal at each of the 6 levels analyzed; n = 7-8 animals/group). In control animals, the number of cells/hilus was greater at ventral levels (Columns 4-6) than at dorsal levels (Columns 1-3). However, the extent of cell loss in pilo-treated animals was greater at all dorsal levels than at ventral levels. There was also greater variability in the extent of cell loss in the ventral dentate hilus.

The qualitative analysis of eYFP-labeled dorsal MCs and CR-labeled ventral MCs, in conjunction with the quantitative analyses of the GluA2-labeled MCs in dorsal and ventral DG, demonstrated a proportionally greater loss of dorsal MCs than ventral MCs in the pilocarpine-treated mice.

### Axonal projections of dorsal MCs were depleted while those of ventral MCs were maintained

The patterns of cell loss led to questions about the axonal projections of MCs in the pilocarpine-treated animals. The axonal projections from dorsal eYFP-labeled MCs and ventral CR-labeled MCs were compared in the same animals in the dorsal and ventral DG. In the dorsal DG of control animals, the CR-labeled fibers formed a distinct, compact band in the inner molecular layer, immediately adjacent to the outer border of the granule cell layer (Fig. 7A). The eYFP-labeled fibers of the dorsal commissural path exhibited a slightly different pattern, with a narrower dense band of fibers located slightly above the granule cell layer, and a more diffuse plexus of fibers that extended into the middle molecular layer (Fig. 7B). When images of dorsal and ventral MC fibers were superimposed, they revealed a trilaminar pattern, with the band of ventral MC fibers located immediately adjacent to the granule cell layer; a dense band of overlapping dorsal and ventral MC fibers in the center; and a diffuse plexus of dorsal MC fibers extending into the middle molecular layer (Fig. 7C).

**Figure 7.**
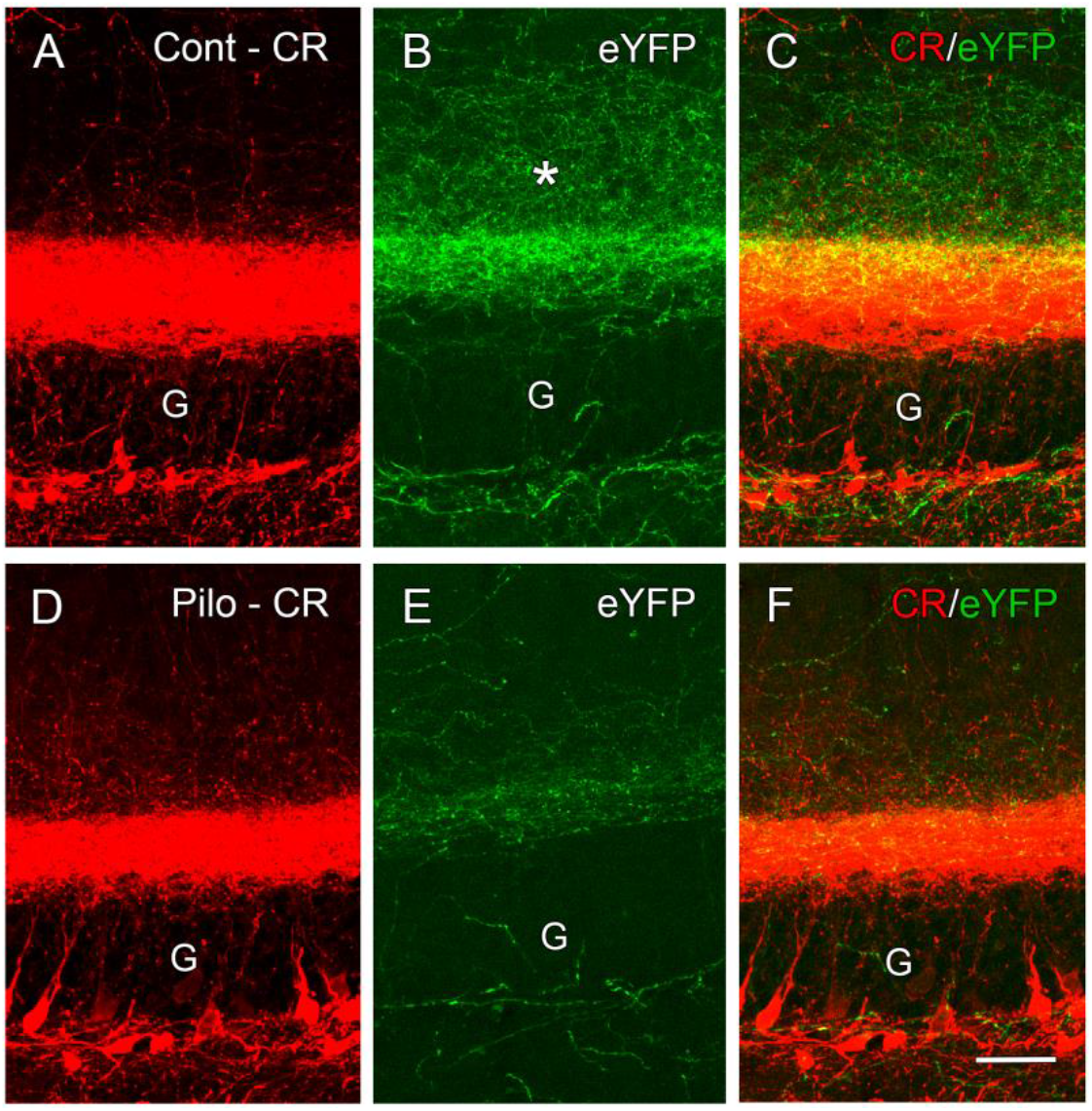
In the dorsal DG, axonal projections from the dorsal and ventral MCs form distinct bands of partially overlapping fibers in the molecular layer of control animals, and this pattern is disrupted in the pilo-treated animals. **A-C**, In control animals, both ventral and dorsal MCs form prominent bands of fibers in the inner molecular layer (**A, B**), but the bands are slightly off-set, with the ventral MC band (**A**) located immediately adjacent to the granule cell layer (G); the dense region of dorsal MC fibers (**B**) located a slight distance away from the granule cell layer; and a diffuse plexus of dorsal MC fibers (*) extending into the middle molecular layer (**B**). This creates a trilaminar pattern of MC innervation (**C**). **D-F**, In pilo-treated animals, the band of ventral MC fibers is relatively well preserved, with some fibers extending beyond the dense band (**D**,**F**) and suggesting some sprouting of remaining ventral MCs. Dorsal MC fibers are severely reduced (**E**), leading to a predominance of ventral MC fibers (**F**). Scale bars: 25 μm (**A-F**).

In the dorsal DG of pilocarpine-treated animals, the ventral MC projection was maintained (Fig. 7D) whereas the dorsal commissural projection was severely depleted (Fig. 7E). The ventral MC projections extended throughout the dorsal DG as a compact band immediately adjacent to the granule cell layer. In contrast, only a limited number of dorsal MC fibers remained, and, when evident, these fibers were in the region of the normally dense portion of the dorsal MC projection (Fig. 7E). The diffuse plexus of fibers in the middle molecular layer was also reduced or absent (Fig. 7E,F).

In the ventral DG, similar decreases in dorsal MC projections and maintained ventral MC projections were observed in the pilocarpine-treated mice (Fig. 3D-F). In horizontal sections, a band of CR labeling was prominent in the inner molecular layer, as in control animals, although there was a decrease in density of labeling compared to the controls (compare Fig. 3A-C with D-F). The dorsal MC projection that normally targets the middle molecular layer at ventral levels (Fig. 3A-C) was severely reduced and often could not be detected in the pilocarpine-treated mice (Fig. 3D-F). The depleted dorsal MC projections and maintained ventral MC projections are consistent with the patterns of GluA2 cell loss in the pilocarpine-treated animals.

The altered patterns of MC fibers appeared to create an imbalance of the dorsal commissural and ventral association pathways throughout the DG in pilocarpine-treated animals, with a predominance of the ventral to dorsal association pathway. The next goal was to determine if there were morphological changes in the remaining MCs that could influence the functions of the pathways.

### Fibers of ventral MCs exhibited sprouting in pilocarpine-treated mice

In control mice, the dense band of ventral MC fibers in the inner one-fourth of the molecular layer had a sharp outer border, and few fibers extended into the middle molecular layer in either the suprapyramidal or infrapyramidal blades of the DG (Fig. 8A,C). In pilocarpine-treated animals, the outer border of the band of ventral MC fibers was more uneven, and labeled fibers emerged from the outer border and extended into the middle molecular layer where they formed a loose plexus of fibers (Fig. 8B). This putative sprouting of the ventral MC axons was observed in both blades of the DG but was more extensive in the infrapyramidal blade (Fig. 8B,D).

**Figure 8.**
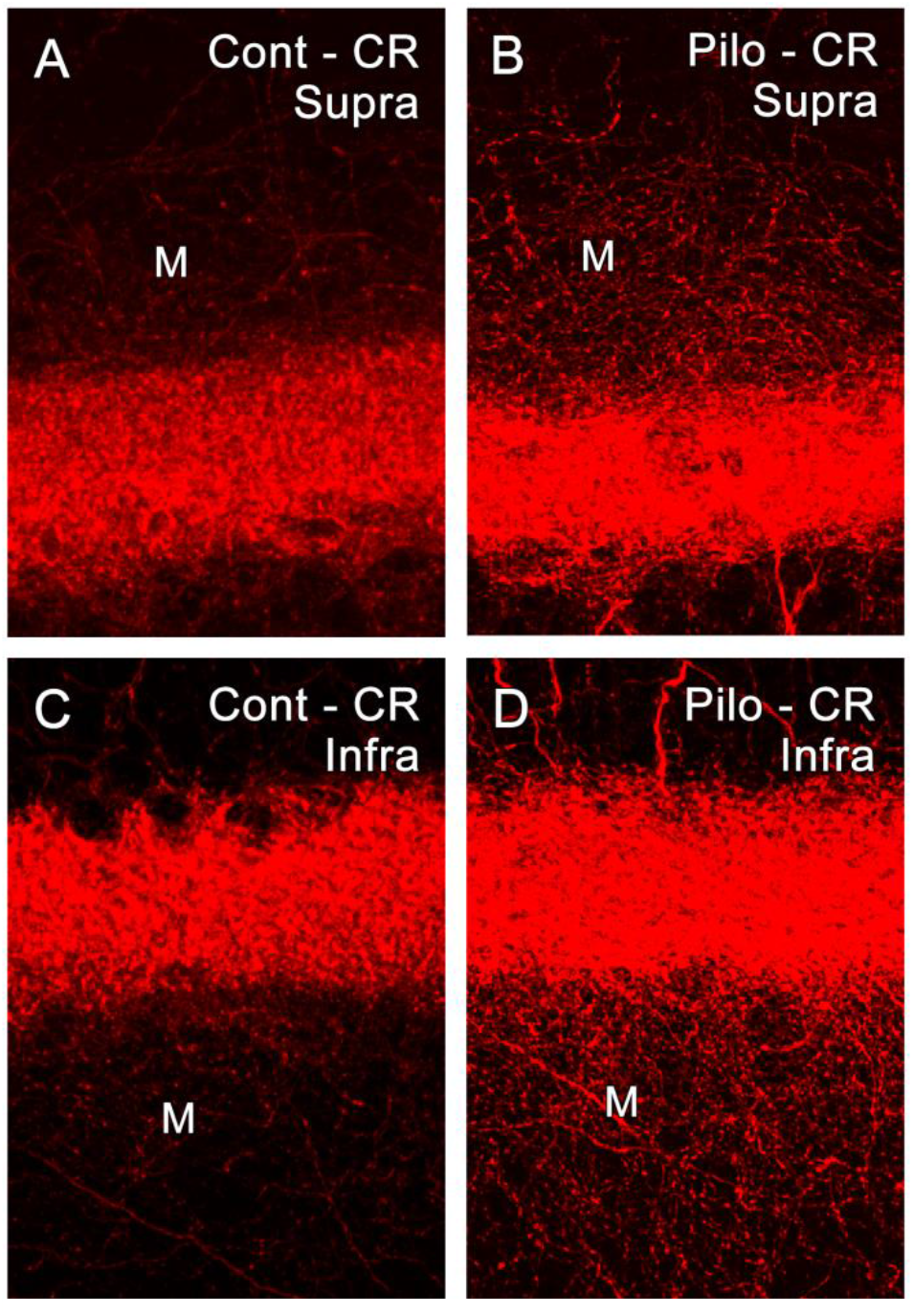
Ventral MC fibers are confined to the inner molecular layer in control animals but extend into the middle molecular layer of the dorsal DG in pilo-treated animals. **A**,**C**, In control animals, the band of CR-labeled fibers from ventral MCs has a relatively sharp outer border, and few labeled fibers extend into the middle molecular layer (M) of the suprapyramidal (**A**) and infrapyramidal (**C**) blades of the DG. **B**,**D**, In pilo-treated animals, the outer border of the CR-labeled MC band is uneven, and a plexus of labeled fibers extends into the middle molecular layer (M) of both blades of the DG. This putative sprouting was often more extensive in the molecular layer of the infrapyramidal blade (**D**). Scale bars: 20 μm (A-D).

We next considered how both loss and preservation of MC fibers could alter the innervation of PV+ interneurons.

### Reduced axonal projections of dorsal MCs in pilocarpine-treated mice could alter innervation of PV-labeled interneurons

Commissural axonal projections from dorsal MCs form synaptic connections with PV+ interneurons as well as hilar GABAergic interneurons in the contralateral DG, and through a disynaptic pathway provide inhibitory control of dentate granule cells. However, the precise sites of termination have been difficult to demonstrate.

In control animals, dendrites of PV interneurons were located directly in the path of the dense band of dorsal MC axons (Fig. 9A), and labeled axonal varicosities were observed in contact with PV+ dendrites in this region (Fig. 9A,C). Considering the electrophysiological evidence for dorsal MC innervation of PV interneurons, this dense band of eYFP-labeled axons would appear to be a likely anatomical site for this innervation. Distal dendritic branches of PV interneurons extended into the more diffuse axonal plexus in the molecular layer, and appositions between the eYFP-labeled varicosities and PV+ dendrites were also observed (Fig. 9A). While the methods in the current study cannot provide conclusive demonstrations of synaptic connections, the direct contacts between MC axonal boutons and dendrites of PV interneurons are consistent with functional connections.

**Figure 9.**
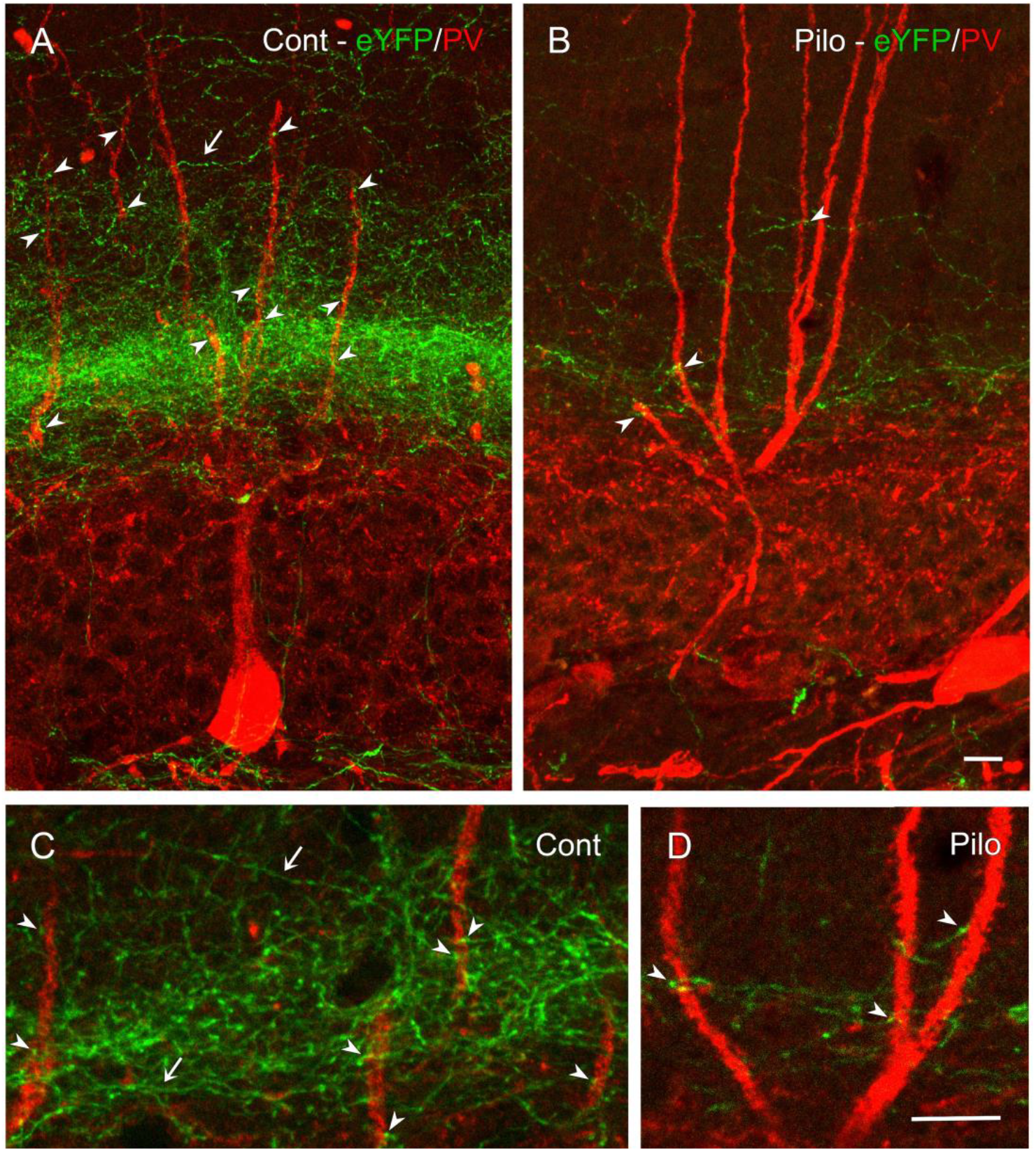
Dendrites of PV+ interneurons extend through the dense plexus of eYFP-labeled fibers of the dorsal commissural path in control animals but are in contact with few eYFP-labeled fibers in pilo-treated animals, due to depletion of the dorsal MC projections. **A**,**C**, The dense band of dorsal MC fibers is in close proximity to dendrites of PV+ interneurons in a control mouse (**A**). Many of these eYFP-labeled fibers extend horizontally through the layer (examples at arrows), and axon terminal-like boutons are adjacent to PV-labeled dendrites (examples at arrowheads (**A**,**C**). **B**,**D**, Such potential contacts (arrowheads) are greatly reduced in pilocarpine-treated animals due to the limited dorsal MC projections (**B**). Scale bars: 10 μm (**A**,**B**); 10 μm (**C**,**D**).

In pilocarpine-treated animals, the axonal plexus of dorsal MCs was severely depleted, and axonal varicosities in contact with dendrites of PV interneurons were similarly reduced (Fig. 9B,D).

### Axonal projections of ventral MCs were in direct contact with dendrites of PV interneurons

In control animals, dendrites of PV+ interneurons were evident within the dense band of ventral MC fibers and terminals (Fig. 10A). However, the high density of CR-labeled fibers and terminals within the band limited the ability to distinguish between contacts of labeled boutons with PV+ dendrites and those with adjacent but unlabeled granule cell dendrites. In pilocarpine treated animals, dendrites of PV+ interneurons were also evident within the band of ventral MC fibers in the inner molecular layer. However, the reduced density of the CR-labeled fibers allowed clearer views of axon terminal-like structures in direct contact with the PV+ dendrites (Fig. 10B), and multiple CR-labeled boutons were evident along the length of the PV+ dendrites in the pilocarpine treated animals (Fig. 10B). These findings suggested an increase or maintenance of ventral MC innervation of PV+ interneurons.

**Figure 10.**
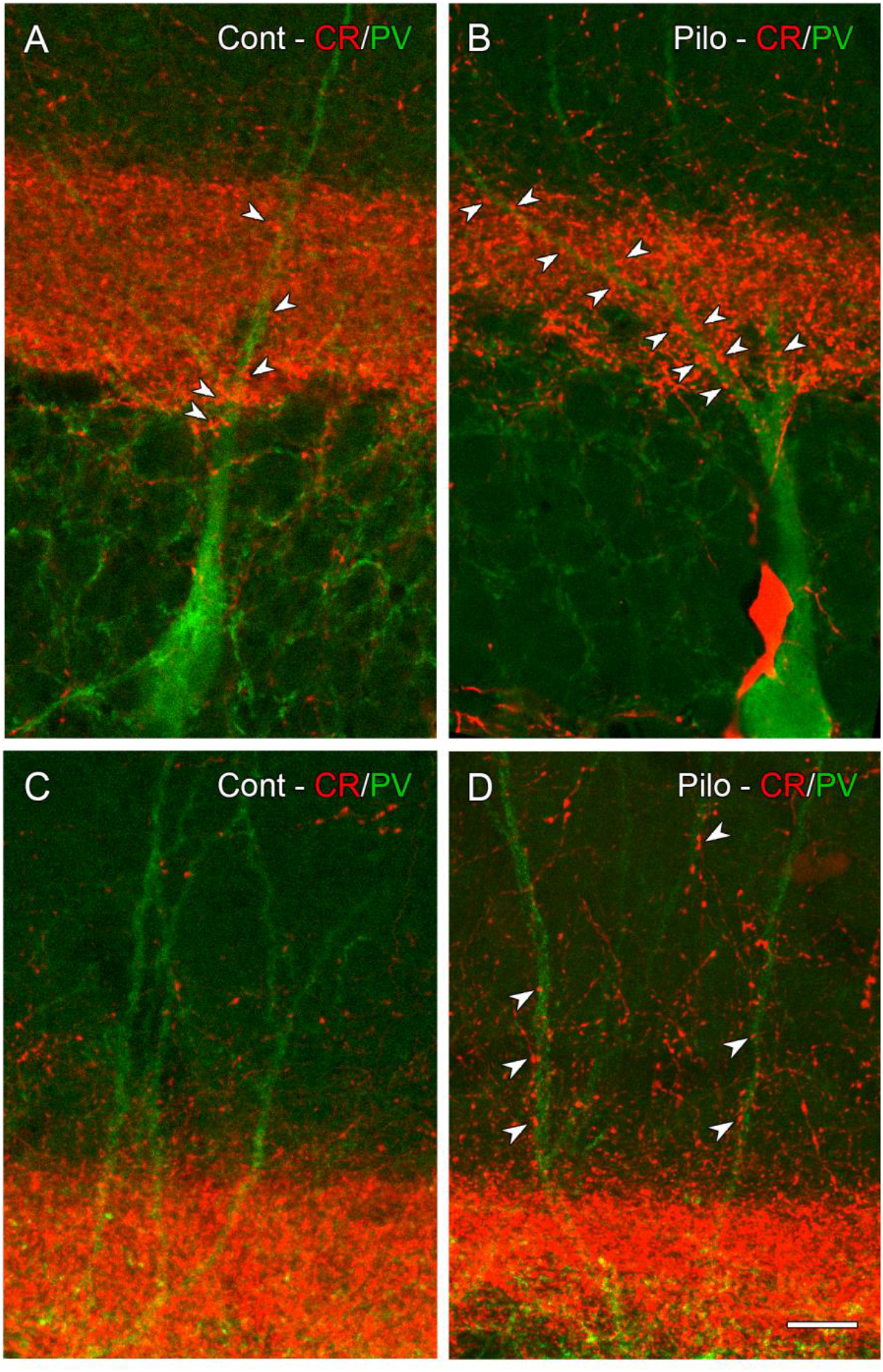
CR-labeled MC fibers are in contact with PV+ dendrites in the inner and middle molecular layer in pilo-treated animals. **A**, In a control animal, some putative contacts between CR-labeled boutons and PV+ dendrites are evident within the dense band of ventral MC fibers (arrowheads), but it is difficult to distinguish between contacts with PV+ dendrites and those with unlabeled granule cell dendrites in the region. **B**, In a pilo-treated animal, a band of ventral MC fibers is present throughout the dorsal DG, as in controls, but appears less dense. Numerous bouton-like structures within the band (examples at arrowheads) are in contact with PV+ dendrites. **C**, In a control animal, only a few CR-labeled fibers extend beyond the dense band of ventral MC fibers, and contacts with PV+ dendrites are limited. **D**, In a pilo-treated animal, putative sprouted CR-labeled fibers extend into the middle molecular layer and multiple labeled boutons are in contact with PV+ dendrites (examples at arrowheads). Scale bar: 10 μm (**A-D**).

PV-labeled dendritic branches could also be identified in the middle molecular layer. In control mice, relatively few CR+ fibers extended into this region and thus few CR+ bouton-like structures were observed in contact with the PV-labeled dendrites (Fig. 10C). In contrast, in pilocarpine-treated animals, CR+ fibers were substantially increased in this region, presumably due to axonal sprouting of ventral MCs, and CR+ boutons were in direct contact with many PV+ dendrites that extended through this region (Fig. 10D).

## Discussion

The major finding of this study is that dorsal and ventral MCs differ in their vulnerability to seizure-induced damage in a mouse model of acquired epilepsy, with severe loss of dorsal MCs and more limited loss of ventral MCs. The dorsal MC loss leads to a marked reduction in the dorsal commissural MC pathway, while the remaining ventral MCs maintain a prominent ventral to dorsal association pathway throughout the DG. These patterns of cell loss create an imbalance between the dorsal and ventral MC pathways that could have functional consequences for the role of MCs in epilepsy. Remaining ventral MCs also exhibit morphological changes that include axonal sprouting and increased or maintained contacts with parvalbumin-labeled interneurons, both of which could affect their function.

### Different patterns of dorsal and ventral MC loss in a mouse model of epilepsy

Both loss and preservation of MCs have been noted in previous studies of epilepsy and TBI models (Lowenstein et al., 1992; Toth et al., 1997; Santhakumar et al., 2000), as well as humans with temporal lobe epilepsy (Blümcke et al., 2000; Seress et al., 2009). These differences in MC fate have been considered broadly and have not been associated with the dorsal and ventral groups of MCs. However, when MCs have been identified by GluA2, a general marker of glutamatergic MCs in the hilus, rostral-caudal differences in MC loss have been noted. A proportionally greater loss of GluA2+ cells in the dorsal DG has been found most consistently in mice and has been described in systemic pilocarpine and kainate models of epilepsy (Volz et al., 2011; Buckmaster et al., 2017). Likewise, a severe loss of GluA2+ cells in the dorsal DG, with considerably less loss in the ventral dentate, has been described in the perforant path stimulation model in mice (Kienzler et al., 2009), and the illustrated patterns of cell loss closely resemble those observed in the current study. Descriptions of cell loss have been more variable in epilepsy models in rats. However, in a detailed study of GluA2+ cell loss following different durations of status epilepticus in the systemic kainate model in rats, more extensive cell loss was found dorsally than ventrally when the duration of status epilepticus was limited to 1-3.5 hrs (Jiao and Nadler, 2007), and this duration is similar to that of the present study. When status epilepticus was allowed to continue for longer durations, a nearly total loss of MCs occurred (Jiao and Nadler, 2007).

It is unclear why dorsal MCs were more vulnerable than ventral MCs in this and previous studies. One possibility is that the CR content of the ventral MCs in the mouse is neuroprotective. Interestingly, MCs in gerbils have high levels of CR expression, in contrast to those in rats which lack CR expression, and, following similar episodes of status epilepticus, CR-containing MCs in the gerbil survived whereas MCs in the rat were severely depleted (Kotti et al., 1996). Also, in a study of MC ablation by diphtheria toxin in the *Calcrl-Cre* mouse, ventral CR-containing MCs degenerated more slowly than dorsal MCs that lack CR expression (Jinde et al., 2012). These findings and those of the present study are consistent with a role for CR in neuroprotection of ventral MCs, although additional factors, including differences in circuitry, could also be involved.

### Imbalance of MC pathways

The loss of dorsal MCs and the preservation of a substantial population of ventral MCs could create an imbalance between two major MC pathways – the dorsal commissural pathway and the ventral to dorsal association pathway. The consequences of these changes will depend on the predominant functions of these pathways, their interactions, and the potential reorganization of remaining MCs. The following discussion is speculative and is based on recent findings in the literature that suggest differences in the function of dorsal and ventral MCs in normal animals. The goal is to suggest a framework for future electrophysiological studies of such differences in animal models of epilepsy.

The major function of MCs has been debated for many years, in part because they form both direct excitatory connections and indirect inhibitory connections with dentate granule cells. Such connections have been demonstrated for all the major MC pathways (Abdulmajeed et al., 2022), and this has led to questions about the predominant function of each pathway *in vivo*.

There is considerable support for a net inhibitory role for the dorsal commissural pathway, observed following activation of dorsal MCs on the contralateral side (Buzsàki and Eidelberg, 1981; Hsu et al., 2016; Danielson et al., 2017). Also, using a computational model of the DG, Danielson et al. (2017) found that deletion of the MC to basket cell connections significantly increased granule cell excitability, whereas deletion of the direct excitatory connections between MCs and granule cells had little effect on granule cell excitability.

While the predominant function of the ventral to dorsal MC pathway continues to be debated (Fredes et al., 2021; Wang et al., 2021), there is increasing support for an excitatory role for this major association pathway. Electrophysiological studies have demonstrated a strong excitation/inhibition balance in this pathway (Abdulmajeed et al., 2022). These findings are supported by behavioral studies in which activation of MCs in the ventral hilus led to increased cFos activity in granule cells of the dorsal dentate (Bauer et al., 2021; Fredes et al., 2021). Electron microscopic studies have also suggested that the synaptic connections of ventral MCs to PV+ interneurons in the dorsal dentate are fewer than those from dorsal MCs (Fredes et al., 2021). Together these findings support a predominantly excitatory role for ventral to dorsal association pathways in normal animals.

Considering these views of the major functions of the commissural and association projections, the loss of dorsal MCs and their inhibitory influences on granule cells, in conjunction with substantial preservation of the ventral MCs and their predominantly excitatory effects, could shift the balance toward increased excitability of dorsal granule cells.

### Plasticity of MC pathways

The imbalance of MC pathways in this mouse model, due to cell loss, would appear to favor a pro-epileptic state, but additional morphological and functional changes could alter the function of remaining MCs.

A recent electrophysiological study of the pilocarpine mouse model of epilepsy demonstrated adaptive changes in the output of remaining MCs, and these led to a preservation or net increase in MC-mediated inhibition of granule cells (Butler et al., 2022). The study did not distinguish between dorsal and ventral MCs, but focused on the intermediate region of the DG where both groups of MCs are likely to be found, and, in this region, both direct and indirect MC connections to granule cells were reduced. However, there was a greater preservation of inhibition, and this enhanced the inhibition to excitation ratio in granule cells. Thus, despite significant cell loss, there was a functionally preserved connection between MCs and PV+ interneurons. The investigators suggested that the MC outputs could reorganize in this epilepsy model and increase their net inhibitory effects in the DG as a homeostatic response to seizure activity.

Several anatomical findings in the current study appear consistent with the electrophysiological findings of Butler et al. (2022). First, the severe loss of dorsal MCs and the decrease in density of the CR+ MC pathway could be related to the described decrease in both excitatory and inhibitory effects of MCs. Also, CR+ boutons were found in contact with PV+ dendrites in the inner molecular layer in the pilocarpine-treated animals, and some of these dendrites appeared to be outlined by labeled terminals. This pattern is consistent with maintained or increased MC innervation of PV+ dendrites in the pilocarpine-treated animals, presumably arising from ventral MCs. These patterns could not be detected in control animals, but this could be due to the normally high density of terminal-like structures in the inner molecular layer and the inability to distinguish between possible contacts on PV+ dendrites and those on adjacent granule cell dendrites.

Sprouting of CR+ fibers was also observed. The CR+ axons and their boutons extended into the middle molecular layer. In this region, CR+ fibers were less dense than in the inner molecular layer, and CR+ boutons could be observed in direct contact with dendrites of PV+ interneurons. Very few CR+ fibers and terminals were observed in this region in control animals, and thus these contacts represent a distinct difference in contacts between MC and PV+ dendrites in the pilocarpine-treated animals and those in controls. If synaptic contacts are formed, this pattern of axonal sprouting could enhance the inhibitory functions of this pathway. These findings are consistent with previous suggestions that an increase in the excitatory innervation of remaining interneurons could partially compensate for impaired inhibition in epilepsy and TBI (Halabisky et al., 2010; Hunt et al., 2011; Butler et al., 2022). While suggestive, the current findings require further comparisons between control and pilocarpine-treated mice and ultrastructural evidence of maintained or increased synaptic contacts of ventral MC terminals on PV-labeled dendrites.

An increase in the inhibitory function of the ventral to dorsal MC association pathway would also be consistent with the findings of Bui et al. (2018) in the intrahippocampal kainate model of epilepsy. Optogenetic activation of the ventral to dorsal MC pathway *in vivo* prevented electrographic seizures from developing into behavioral seizures. This led to the suggestion that surviving ventral MCs have a net inhibitory effect and play a protective role in preventing spontaneous seizure progression. This study was conducted prior to descriptions of distinct populations of dorsal and ventral MCs in the mouse, and it is possible that the stimulation was primarily in the intermediate region of the DG, where both dorsal and ventral MCs could have been activated (Fredes et al., 2021). However, if ventral MC fibers were selectively stimulated, the findings could suggest a shift from a predominantly excitatory to an inhibitory role of this association pathway.

Previously reported changes in remaining ventral MCs could also alter their function. These include increases in somal area and mEPSP frequency, which are consistent with increased excitatory input to remaining MCs (Zhang et al., 2015). However, the overall functional effect of these MC changes would depend on the predominant targets of these ventral MCs – whether granule cells or GABAergic interneurons.

While morphological changes were the focus of this study, the function of remaining MCs could also be modified in response to different patterns and strength of stimulation. Previous studies of the dorsal DG have demonstrated that the normal, predominantly inhibitory, commissural pathway can be transformed into a excitatory pathway through stimulation that elicits a unique form of mossy cell to granule cell long-term potentiation (LTP), mediated by brain-derived neurotrophic factor (BDNF) (Hashimotodani et al., 2017). In recent studies of seizure induction in epilepsy models, optogenetic stimulation of MCs, utilizing the previously identified stimulation patterns, strengthened seizure activity following the systemic administration of kainate, suggesting that such stimulation was proconvulsant (Nasrallah et al., 2022). Likewise, strong stimulation of MCs, as occurs during induction of status epilepticus by pilocarpine, can lead to powerful excitation of granule cells (Botterill et al., 2019). These findings are consistent with an abundance of excitatory MC contacts on granule cell dendrites (Buckmaster et al., 1996; Wenzel et al., 1997).

### Hypothesized roles of MCs in epilepsy

Previous studies of MCs in epilepsy models have led to two different hypotheses. Both suggest that MCs could contribute to increased excitability and a pro-epileptic state, although by different processes. The “dormant basket cell hypothesis” emphasizes the loss of MCs and associated loss of their innervation of GABAergic basket cells, leading to increased excitability and seizure susceptibility within the DG (Sloviter, 1991; Sloviter et al., 2003). In contrast, the “irritable MC hypothesis” focuses on the remaining MCs and suggests that increased activity of these cells could increase excitation throughout the DG (Santhakumar et al., 2000; Ratzliff et al., 2002). Interestingly, the extensive loss of MCs in the dorsal DG in the present study appears most consistent with the dormant basket cell hypothesis whereas the relative preservation of ventral MCs would be most consistent with the irritable MC hypothesis.

The patterns of MC loss, when considered in relation to the potentially different roles of dorsal and ventral MCs, could lead to a broader view of the function of MCs in epilepsy. Likewise, recent findings of adaptive changes in MC circuitry in an epilepsy model add a new dimension to the role of remaining MCs in epilepsy (Butler et al., 2022).

### Limitations of the study

The patterns of MC loss observed in the present study are unlikely to be found in all mouse models of acquired epilepsy and TBI, and in other species with different patterns of CR expression in MCs. Even in the same model, the strength and duration of status epilepticus can alter the patterns of MC loss. Nevertheless, differences in the extent of dorsal and ventral MC loss are important to consider as they could influence the seizure initiation sites and the propagation of seizure activity throughout the DG.

The current study focused on patterns and interactions of two major MC pathways – the dorsal commissural and the ventral to dorsal association pathways. However, the loss of other MC pathways, including the dorsal to ventral association pathways and potential alterations in commissural connections of ventral MCs, could also influence seizure activity.

This study has emphasized different patterns of MC loss and the potential plasticity of remaining MCs. Numerous other changes in the DG, including loss of hilar interneurons and reorganization of mossy fibers of dentate granule cells, are occurring simultaneously, and these may interact with and be influenced by the alterations in MCs.

### Continued challenges

By differentially labeling dorsal and ventral MCs, we were able to identify distinct differences in the axonal projections of these MCs in an epilepsy model. The observed imbalance between the dorsal and ventral MC systems in this model raises questions about the normal interactions of these pathways and the consequences of their imbalance in epilepsy. The findings re-emphasize the importance of the ventral to dorsal MC association pathway in epilepsy and the need to study the function of this pathway *in vivo*.

